# Streptococcal dTDP-L-rhamnose biosynthesis enzymes: functional characterization and lead compound identification

**DOI:** 10.1101/312157

**Authors:** Samantha L. van der Beek, Azul Zorzoli, Ebru Çanak, Robert N. Chapman, Benjamin H. Meyer, Geert-Jan Boons, Helge C. Dorfmueller, Nina M. van Sorge

## Abstract

Biosynthesis of the nucleotide sugar precursor dTDP-L-rhamnose is critical for the viability and virulence of many human pathogenic bacteria, including *Streptococcus pyogenes* (Group A *Streptococcus*; GAS) and *Streptococcus mutans*. Those pathogens require dTDP-L-rhamnose for the production of structurally similar rhamnose polysaccharides in their cell wall. Via heterologous expression in *S. mutans*, we confirm that GAS RmlB and RmlC are critical for dTDP-L-rhamnose biosynthesis through their action as dTDP-glucose-4,6-dehydratase and dTDP-4-keto-6-deoxyglucose-3,5-epimerase enzymes, respectively. Complementation with GAS RmlB and RmlC containing specific point mutations corroborated the conservation of previous identified catalytic residues in these enzymes. Bio-layer interferometry was used to identify and confirm inhibitory lead compounds that bind to GAS dTDP-rhamnose biosynthesis enzymes RmlB, RmlC and GacA. One of the identified compounds, Ri03, inhibited growth of GAS as well as several other rhamnose-dependent streptococcal pathogens with an MIC_50_ of 120-410 μM. We therefore conclude that inhibition of dTDP-L-rhamnose biosynthesis such as Ri03 affect streptococcal viability and can serve as a lead compound for the development of a new class of antibiotics that targets dTDP-rhamnose biosynthesis in pathogenic bacteria.

## Introduction

The typical cell wall architecture of a Gram-positive bacterium consists of a thick peptidoglycan layer that is decorated with wall teichoic acids, proteins and capsular polysaccharides. However, certain lactic acid bacteria, particularly streptococcal species, lack the expression of wall teichoic acids and instead express rhamnose cell wall polysaccharides, which are covalently anchored to peptidoglycan (Mistou *et al.*, 2016). Rhamnose cell wall polysaccharides encompass about 40-60% of the cell wall mass and are considered to be functional homologs of wall teichoic acids (Caliot *et al.*, 2012, Sutcliffe *et al.*, 2008). Examples of rhamnose cell wall polysaccharides are the group A carbohydrate (GAC) of Group A *Streptococcus* (GAS; *Streptococcus pyogenes*) and the serotype-determining polysaccharides, referred to as rhamnose-glucose polysaccharides (RGP), of *Streptococcus mutans*. The GAC and RGP share structural similarities; both consist of an α-1,2-/α-1,3-linked polyrhamnose backbone with alternating *N*-acetylglucosamine side chains for GAS and glucose or galactose side chains for *S. mutans* (Heymann *et al.*, 1964, Huang *et al.*, 1986, St Michael *et al.*, 2017, Pritchard *et al.*, 1986, Linzer *et al.*, 1987) (Fig. 1). For *S. mutans*, the type of side chain as well as their linkage to the rhamnan backbone determines the *S. mutans* serotype (Nakano & Ooshima, 2009, St Michael *et al.*, 2017) (Fig. 1). In contrast, all GAS serotypes express a structurally invariant GAC (Lancefield, 1933). Similar to classical wall teichoic acids, rhamnose cell wall polysaccharides are critical for maintaining cell wall shape, bacterial physiology and virulence, but in-depth knowledge of their biosynthesis or host interactions at a molecular level is limited (van der Beek *et al.*, 2015, van Sorge *et al.*, 2014, Kovacs *et al.*, 2017, Zhu *et al.*, 2017). A better understanding of these mechanisms could aid the development of new classes of antibiotics, antibiotic adjuvants or vaccines (St Michael *et al.*, 2017, van Sorge *et al.*, 2014, Sabharwal *et al.*, 2006).

**Fig. 1.**
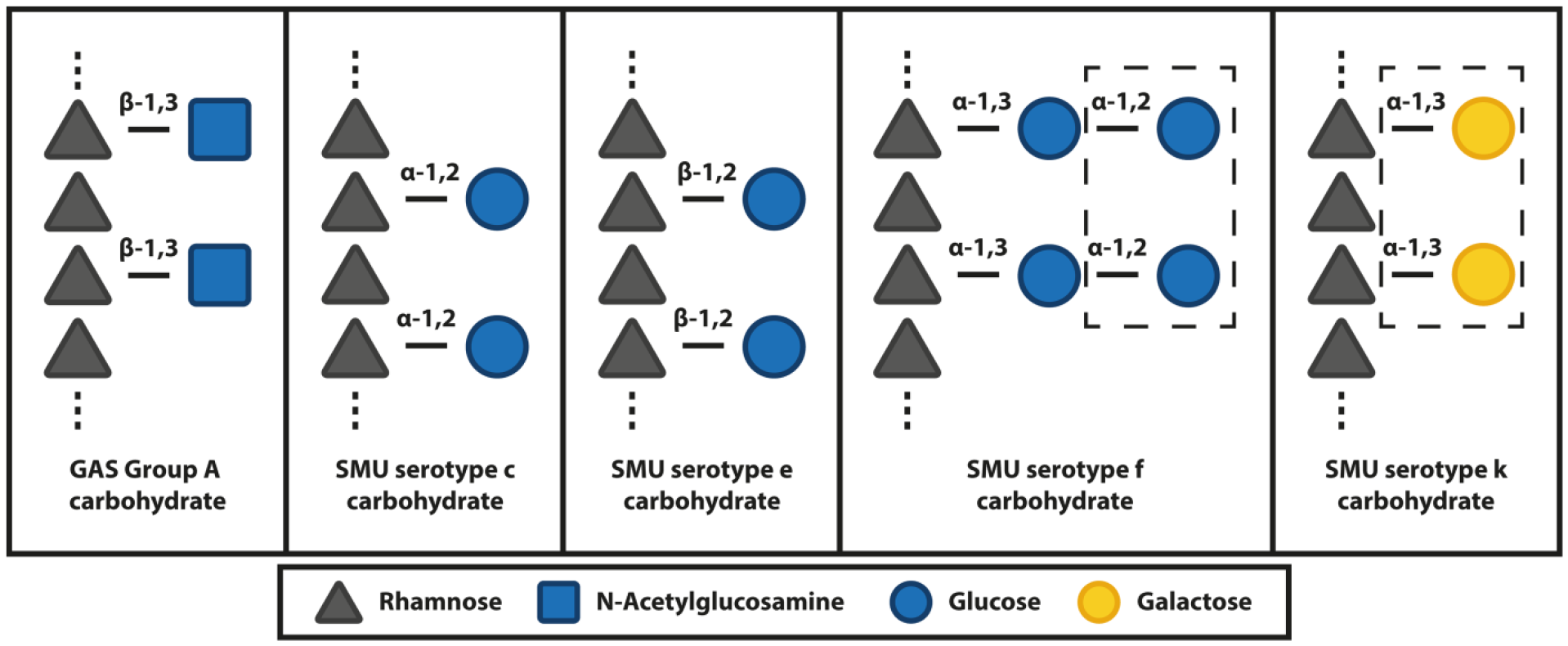
Schematic overview of streptococcal rhamnose polysaccharide structures. Schematic representation of the chemical composition of the group A carbohydrate from GAS and the serotype c, e, f and k carbohydrates from *S. mutans* (SMU). All carbohydrates share an α-1,2/α-1,3 linked polyrhamnose backbone. Sugar residues in dashed boxes were recently identified by St Michael *et al.* (6).

L-Rhamnose is the main building block for both the GAC and RGP. The biosynthesis pathway of the nucleotide precursor, dTDP-L-rhamnose, is highly conserved among both Gram-positive and Gram-negative bacteria (Mistou *et al.*, 2016, Giraud & Naismith, 2000). The dTDP-L-rhamnose biosynthesis pathway is critical or even essential for viability or virulence of a wide range of human pathogens including GAS (Le Breton *et al.*., 2015, Le Breton *et al.*, 2013), Group B *Streptococcus* (GBS) (Hooven *et al.*, 2016, Caliot *et al.*, 2012), some serotypes of *Streptococcus pneumoniae* (Kim *et al.*, 1999, Magee & Yother, 2001), *S. mutans* (Engels-Deutsch *et al.*, 2003, Kovacs *et al.*, 2017, Shields *et al.*, 2018, Tsuda *et al.*, 2000), *Enterococcus faecalis* (Xu *et al.*, 2000, Rigottier-Gois *et al.*, 2015), *Mycobacterium spp*. (Li *et al.*, 2006, Ma *et al.*, 2002), *Pseudomonas spp*. (Engels *et al.*, 1985) and *Salmonella enterica* serovar *Typhimurium* (Joiner, 1988). Consequently, this pathway is considered to be an interesting drug target, especially since dTDP-L-rhamnose is not produced or used by humans (Adibekian *et al.*, 2011). dTDP-L-rhamnose is produced through a four-step enzymatic pathway catalyzed by the enzymes RmlABCD. In the first step of the pathway, RmlA, a glucose-1-phosphate thymidyltransferase, converts glucose-l-phosphate into dTDP-glucose (Blankenfeldt *et al.*, 2000), which is subsequently oxidized and dehydrated to form dTDP-4-keto-6-deoxy-D-glucose by the dTDP-D-glucose 4,6-dehydratase RmlB (Beis *et al.*, 2003). RmlC catalyzes an unusual double epimerization reaction (Dong *et al.*, 2007, Giraud *et al.*, 2000, Dong *et al.*, 2003b), the product of which is finally reduced by RmlD, a dTDP-4-dehydrorhamnosereductase, to form dTDP-L-rhamnose (Blankenfeldt *et al.*, 2002, van der Beek *et al.*, 2015). The mechanisms of action as well as structural characteristics of the RmlABCD enzymes have been studied extensively (Dong *et al.*, 2003a, Dong *et al.*, 2003b, van der Beek *et al.*, 2015, Giraud *et al.*, 2000, Beis *et al.*, 2003, Kantardjieff *et al.*, 2004, Allard *et al.*, 2001, Allard *et al.*, 2002, Dong *et al.*, 2007, Blankenfeldt *et al.*, 2000). Indeed, crystal structures are available for all four Rml enzymes from different Gram-positive and Gram-negative bacterial species and show high structural conservation. This structural and functional information has enabled the development of different screening methods to discover inhibitors against RmlBCD, yielding compounds that can inhibit dTDP-L-rhamnose biosynthesis in the low micromolar range in a biochemical assay (Babaoglu *et al.*, 2003, Ma *et al.*, 2001, Ren *et al.*, 2015, Sivendran *et al.*, 2010, Wang *et al.*, 2011). Given the importance of L-rhamnose to virulence and viability of many human bacterial pathogens, potent inhibitors of these pathways are of therapeutic interest, especially with the alarming development of antibiotic resistant bacteria.

Recently, we characterized GAS GacA as an RmlD homolog catalyzing the last step in dTDP-L-rhamnose biosynthesis (van der Beek *et al.*, 2015). Surprisingly, this GAS RmlD enzyme was confirmed to be a monomer as opposed to previously characterized RmlD dimers. Additional bioinformatics analysis of 213 RmlD homologs revealed that the majority of RmlD enzymes are predicted to be monomeric, while only a subclass of RmlD enzymes in Gram-negatives form metal-dependent homodimers (Blankenfeldt *et al.*, 2002). In this study, we extended our work to study GAS RmlB and RmlC on a functional level through heterologous expression in *S. mutans* and subsequent analysis of growth, morphology and cell wall composition. In addition, we report the identification of small chemical fragments that bind these enzymes and inhibit GAS growth in a sub- to low-millimolar range. Furthermore, the most potent fragment, Ri03, could inhibit the growth of *S. mutans* and *S. equi* subsp. *zooepidemicus* (Group C *Streptococcus*, GCS) with similar efficacy. These results demonstrate that rhamnose biosynthesis inhibitors can directly interfere with bacterial viability and could form a new class of antibiotics targeting nucleotide sugar production.

## Results

### Protein sequence analysis of GAS RmlB and RmlC

As an extension of our previous work on GAS GacA (van der Beek *et al.*, 2015), we sought to characterize GAS RmlB (GAS5448_RS05645) and RmlC (GAS5448_RS05650), the putative dTDP-glucose-4,6-dehydratase and dTDP-4-keto-6-deoxyglucose-3,5-epimerase, respectively. GAS *rmlB* and *rmlC* are clustered in an operon together with *rmlA* in the order *rmlACB*. Structural and biochemical analysis of RmlB and RmlC from *S. suis*, *S. enterica* and *M. tuberculosis* revealed that both enzymes are functional homodimers in these organisms (Beis *et al.*, 2003, Dong *et al.*, 2003b, Allard *et al.*, 2001, Giraud *et al.*, 2000, Kantardjieff *et al.*, 2004). Protein sequence alignment of RmlB and RmlC homologs from several streptococcal species and *S. enterica* displayed high homology (Fig. 2; Table S2). Importantly, all catalytic residues in RmlB and RmlC are conserved (Fig. 2; Table S2).

**Fig. 2.**
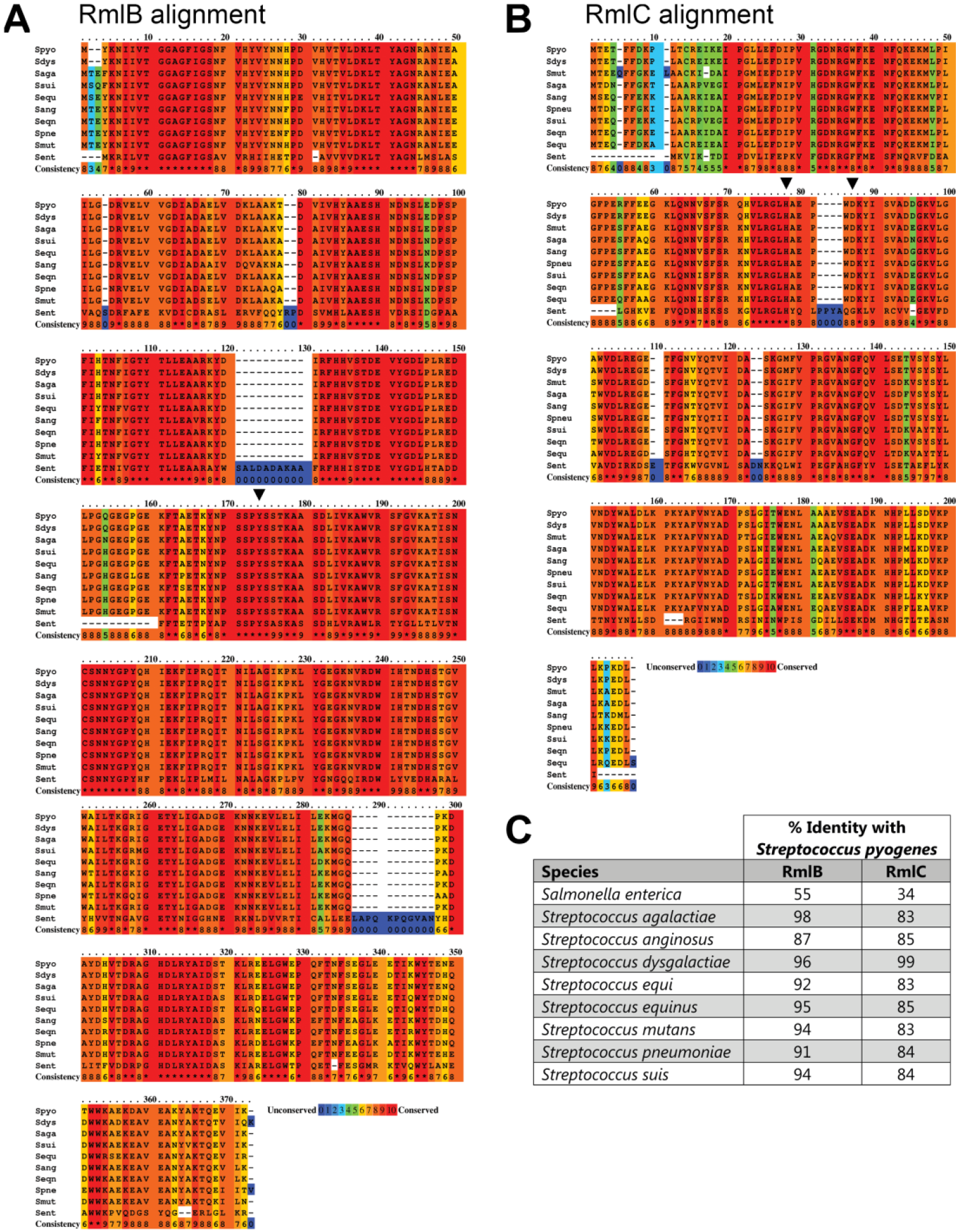
Protein sequence alignment and identity matrix of RmlB and RmlC homologs. Color-coded representation of amino acid conservation for (A) GAS RmlB and (B) GAS RmlC to *S. enterica* and other streptococcal species. The amino acid conservation is scored from 0 to 10, with 0 (color blue) assigned to the least conserved residue and 10 (color red) to the most conserved residue. Critical enzymatic residues for RmlB (Y159) and RmlC (H76 and K82) are indicated with an inverted triangle. *S. pyogenes* (Spyo); *S. enterica*; (Sent); *S. agalactiae* (Saga); *Streptococcus anginosus* (Sang); *Streptococcus dysgalactiae* (Sdys); *Streptococcus equi* (Sequ); *Streptococcus equinus* (Seqn); *S. mutans* (Smut); *S. pneumoniae* (Spneu); *Streptococcus suis* (Ssui). Protein accession numbers are described in the supplementary data (Table S2). (C) Percentage identity matrix of RmlB and RmlC homologs.

### GAS RmlB and RmlC functionally replace *S. mutans* homologs

Most genes directly or indirectly involved in the GAC biosynthesis pathway are essential for GAS viability, including all four dTDP-L-rhamnose biosynthesis genes *rmlABC* and *gacA* (Le Breton *et al.*, 2015, Le Breton *et al.*, 2013, van der Beek *et al.*, 2015, Zhu *et al.*, 2017). In *S. mutans*, the dTDP-L-rhamnose biosynthesis genes are not essential under non-competitive conditions, although gene deletions result in attenuated growth and severe morphological defects (van der Beek *et al.*, 2015, Tsukioka *et al.*, 1997). GAS RmlB and RmlC proteins share 94% and 83% protein sequence identity with the *S. mutans* homologs, respectively. In addition, the organization of these genes is identical in both species with the exception of a hypothetical protein (174 nucleotides, 57 amino acids) encoded between RmlC and RmlB.

To confirm the function of GAS RmlB and RmlC in dTDP-L-rhamnose biosynthesis, we heterologously expressed GAS RmlB and GAS RmlC-encoding genes in *S. mutans* strains lacking *rmlB* or *rmlC*, respectively. Deletion of *S. mutans rmlB* (SMU Δ*rmlB*) or *rmlC* (SMU Δ*rmlC*) by replacement with an erythromycin resistance cassette severely attenuated bacterial growth compared to the wild-type strain (WT) (Fig. 3), which is characteristic for a rhamnose-deficient *S. mutans* strain and in line with our previously constructed *S. mutans rmlD* deletion strain (van der Beek *et al.*, 2015). Morphological analysis of *rmlB* and *rmlC* mutant bacteria by scanning electron microscopy revealed swelling and clumping of bacteria as a result of misplaced septa resulting in division errors and multidirectional growth (Fig. 4A). Subsequent analysis of the cell wall carbohydrate composition by HPLC/MS confirmed that SMU Δ*rmlB* and SMU Δ*rmlC* lacked rhamnose in their cell walls, which concordantly resulted in the loss of the glucose side chains (Fig. 4B). Introduction of either homologous *S. mutans rmlB* or heterologous GAS *rmlB* on an expression plasmid in the corresponding SMU Δ*rmlB* mutant restored rhamnose incorporation in the cell wall (Fig. 4B) as well as the defective morphological phenotype and growth (Fig. 3, 4A). Initially, we were unable to complement SMU Δ*rmlC* with *rmlC* from *S. mutans*, whereas heterologous GAS *rmlC* could restore growth, morphology and rhamnose production of the SMU Δ*rmlC* mutant (Fig. 3B, 4). Upon reexamination of the UA159 genome, a *S. mutans* serotype c strain that we used as a reference genome for *S. mutans* Xc, we located an alternative ATG-start site 135 bp upstream of the annotated *rmlC* gene, which is in line with the annotation of *rmlC* in GAS. Importantly, available structural information indicates that the first 45 amino acids are part of the RmlC dimerization interface, in particular forming the extension of the beta-sheet with two additional beta-strands, which are required for nucleotide binding (Christendat *et al.*, 2000, Giraud *et al.*, 2000). In agreement with structural and genetic predictions, complementation of SMU Δ*rmlC* with the extended PCR product complemented all observed defects (Fig 3B, 4), indicating that the extended genomic PCR product encodes a functional RmlC enzyme (198 amino acids), which is similar to the size of GAS and *S. suis* RmlC (197 amino acids).

**Fig. 3.**
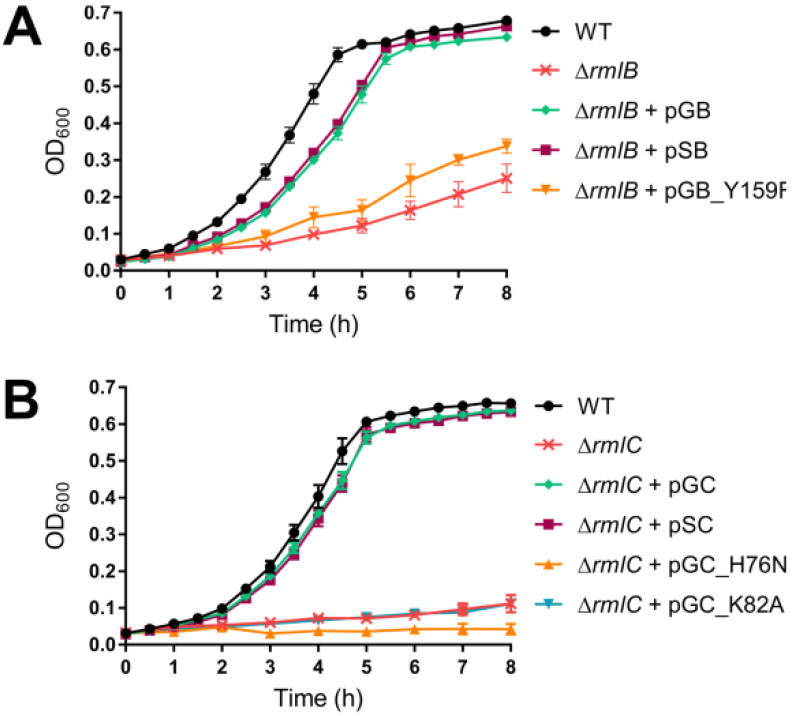
Heterologous expression of GAS RmlB and RmlC and catalytically-inactive enzymes in S. mutans: growth curves. Growth curves of *S. mutans* (A) *rmlB* and (B) *rmlC* mutant sets: wild-type (WT), Δ*rmlB*, Δ*rmlC*, Δ*rmlB* + pSB, Δ*rmlB* + pGB(_Y159F), Δ*rmlC* + pSC and Δ*rmlC* + pGC(_H76N/K82A). Growth curves represent mean ± standard error of mean (SEM) of at least three biological repeats.

**Fig. 4.**
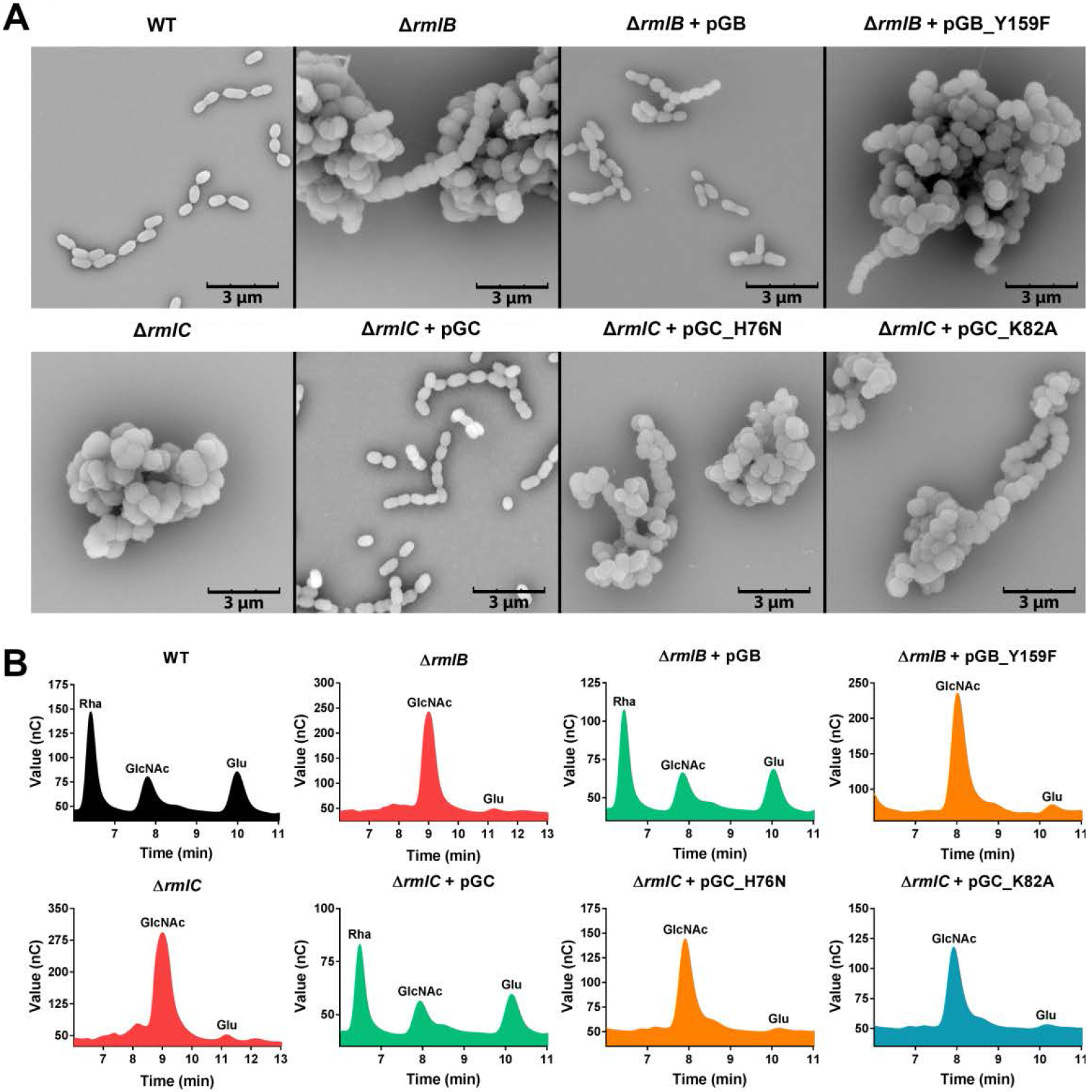
Heterologous expression of GAS RmlB and RmlC and catalytically-inactive enzymes in *S. mutans*: morphology and cell wall content. (A) Representative scanning electron microscopy images and (B) cell wall carbohydrate composition analysis of *S. mutans* wild-type (WT), Δ*rmlB*, Δ*rmlC*, Δ*rmlB* + pSB, Δ*rmlB* + pGB(_Y159F), Δ*rmlC* + pSC and Δ*rmlC* + pGC(_H76N/K82A). Rha, rhamnose; GlcNAc, N-acetylglucosamine; Glu, glucose.

### Catalytic residues of GAS RmlB and RmlC are conserved

Based on biochemical conformation of catalytic residues in *S. suis* RmlB and RmlC (Allard *et al.*, 2001, Dong *et al.*., 2003b) and protein sequence alignment with the GAS RmlB and RmlC homologs (Fig. 2), we set out to functionally validate predicted catalytic residues of GAS RmlB (Y159) and RmlC (H76 and K82). Point mutations were individually introduced in GAS RmlB (Y159F) and RmlC (H76N and K82A) and overexpression vectors carrying these mutant genes were expressed in SMU Δ*rmlB* and SMU Δ*rmlC*, respectively. All mutated residues are involved in catalysis and mutated genes were unable to complement the characteristic rhamnose-depleted phenotype consisting of growth retardation and an aberrant morphology (Fig. 3, 4A). Complementary, our cell wall composition analysis revealed that the *S. mutans* strains carrying these three mutated constructs completely lacked incorporation of rhamnose in their cell wall (Fig. 4B).

### Inhibitor screen against GAS Rml proteins and hit confirmation

Given the importance of the dTDP-L-rhamnose biosynthesis pathway for viability or virulence of many human pathogens, we aimed to identify chemical scaffolds that could act as starting points for future optimization and drug development pipelines. Therefore, we conducted a bio-layer interferometry (BLI) inhibitor screen against the commercially available Maybridge Library using the three recombinant GAS dTDP-rhamnose biosynthesis enzymes RmlB, RmlC and GacA (Fig. S1A). The advantage of this approach is that it precludes the use of expensive or commercially unavailable enzyme substrates. Using a commercially available library of ^~^1,000 chemical fragments, our initial screen identified 12 hits (Fig. S1B-D) of which seven fragments were found to specifically interact with GAS RmlB/RmlC/GacA as confirmed by BLI (Fig. 5, S2). In addition, we confirmed the identified hits in dilution series for their binding affinity to these three enzymes (Fig. S2, Table S3). Importantly, this method allowed us to estimate enzyme specificity by comparing the data obtained from one enzyme to the other two. Using this strategy, we identified chemical scaffolds with specificity for one enzyme, but also several fragments that bound to more than one enzyme of the dTDP-rhamnose biosynthesis pathway (Fig. 5, S2, Table S3).

**Fig. 5.**
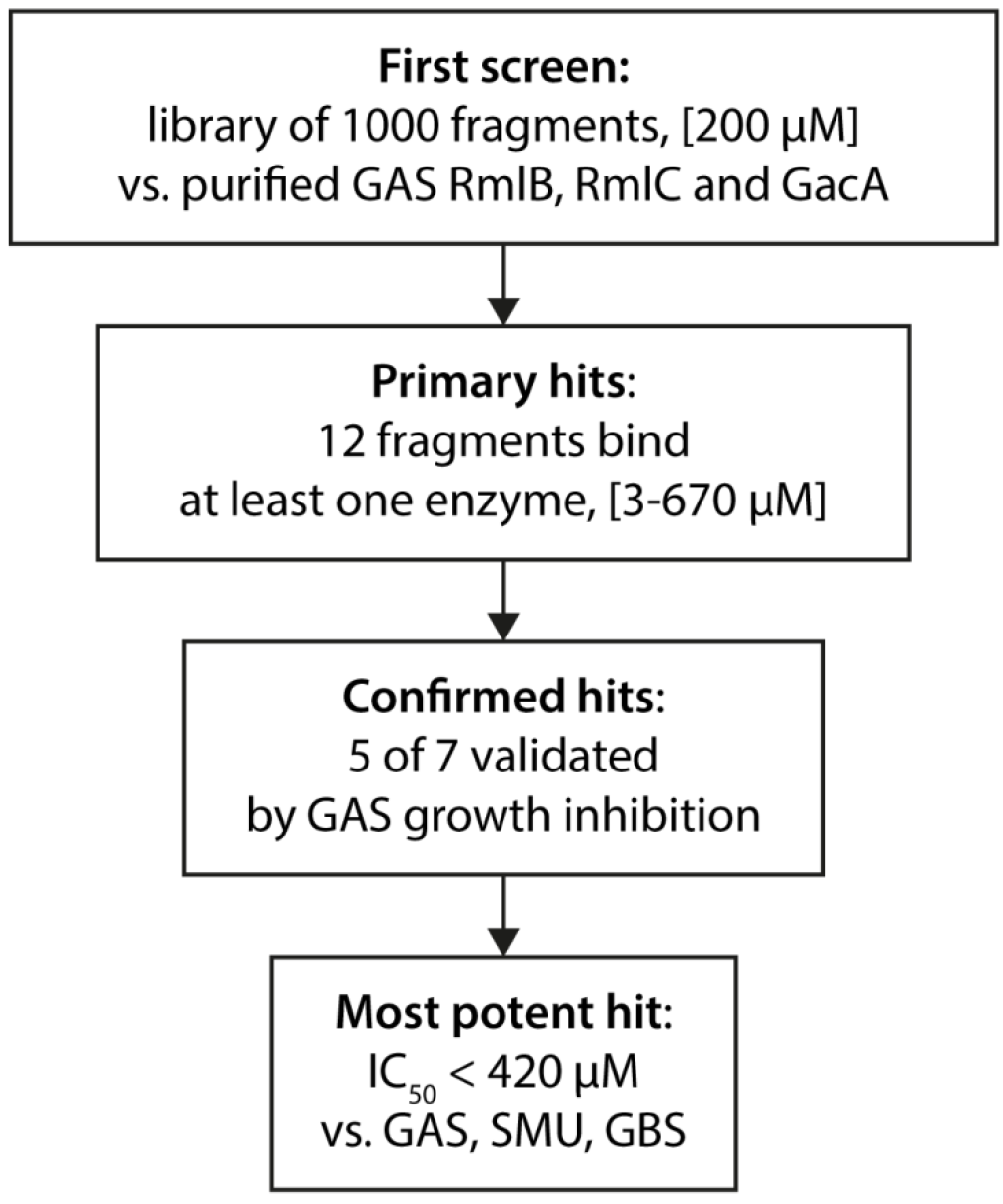
Screening and fragment validation flowchart.

### Bactericidal activity of identified fragments

Since dTDP-rhamnose is an essential nucleotide sugar in GAS (Le Breton *et al.*, 2017, Le Breton *et al.*, 2015), we assessed the functional capacity of these scaffolds to impact GAS growth. All fragments identified in this study were water insoluble and were therefore dissolved in DMSO, except Ri07, which was already liquid at room temperature. GAS is tolerant to DMSO concentrations of 2%, therefore this was the maximum concentration used in bacterial assays. Even in DMSO, Ri04 and Ri07 were highly insoluble in bacterial culture medium and could therefore not be tested for inhibition of bacterial growth. The remaining five compounds were soluble at the highest concentration tested and were able to inhibit growth of GAS with IC_50_ values ranging from 110 μM to 6. 2 mM (Fig. 5, 6A, Table 1). In accordance with growth inhibition, GAS morphology was also severely affected upon exposure to either 200 μM Ri03 or 100 μM Ri06 (Fig. 6C). Especially for Ri03, streptococci were swollen and chains were longer, a phenotype reminiscent of an inducible *gacA* GAS knock-out strain (van der Beek *et al.*, 2015).

**Fig 6.**
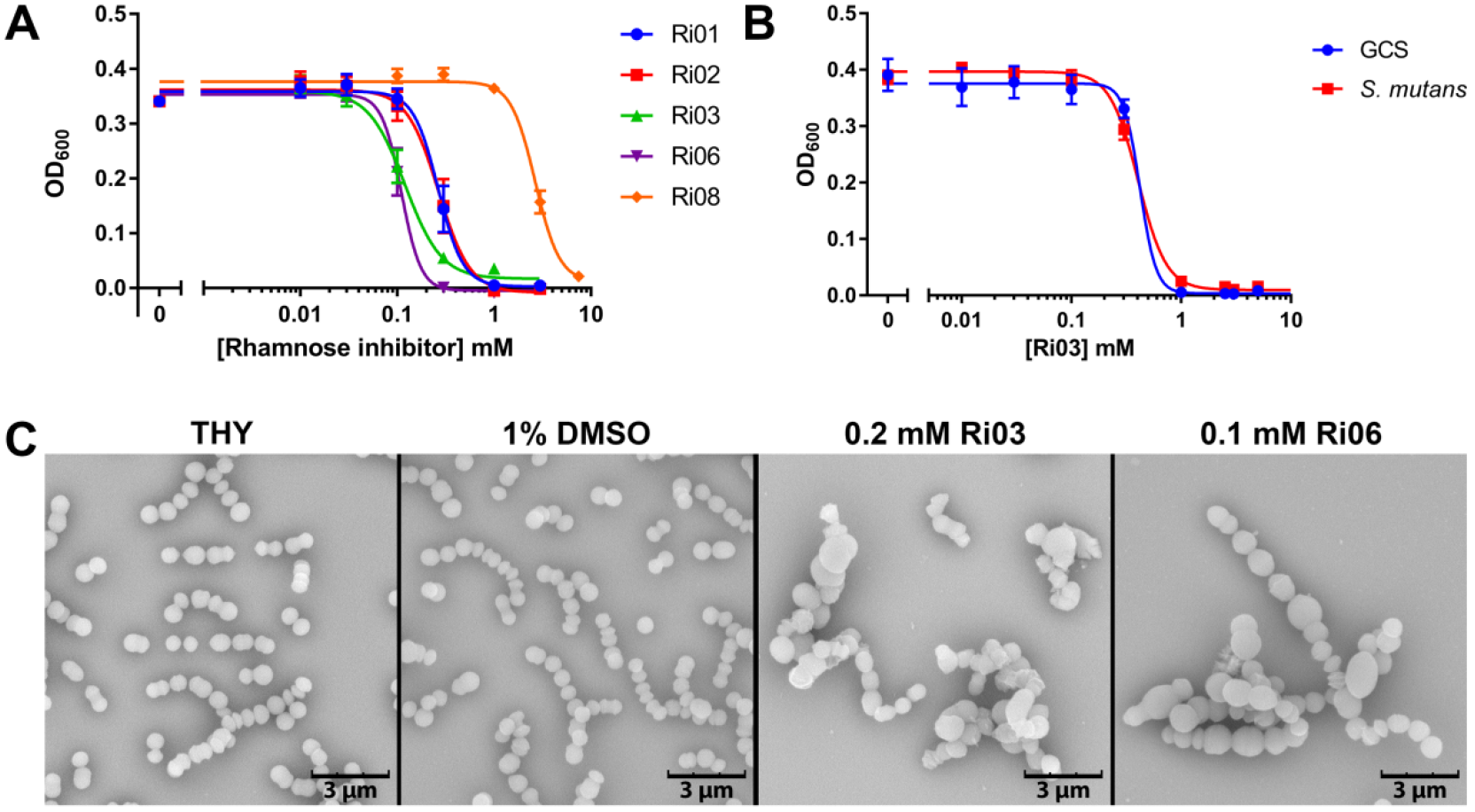
Identified fragments inhibit growth of streptococcal species. (A) Dose-response curves for growth inhibition of GAS with various concentrations of Ri01, Ri02, Ri03, Ri06or RiOS. IC_50_ values rangefrom 110 μM to 2.66 mM. (B) Dose-response curves for growth inhibition of *S. equi* subsp. *zooepidemicus* (Group C *Streptococcus*; GCS) and *S. mutans* with various concentrations of Ri03. IC_50_ values are 420 μM and 410 μM, respectively. Dose-response curves represent mean ± SEM of at least three biological experiments performed in duplicate. (C) Representative scanning electron microscopy images of GAS after 16 h incubation in growth medium (THY), or THY with the addition of 1% DMSO, 0.2 mM Ri03 or 0.1 mM Ri06.

**Table 1.** Overview of identified compounds and IC_50_ for GAS growth inhibition.

| Inhibitor | Compound | Binding target | IC<sub>50</sub> (mM) GAS |
| --- | --- | --- | --- |
| Ri01 | N-(1-Naphthyl)ethylenediamine dihydrochloride | RmlB + RmlC | 0.27 |
| Ri02 | 2,5-Difluorophenylhydrazine | RmlB + GacA | 0.27 |
| Ri03 | 5-(4-chlorophenyl)-2-furoic acid | RmlB + GacA | 0.12 |
| Ri06 | 4,4'-Thiodiphenol | RmlB + GacA | 0.11 |
| Ri08 | 2-(2,5-Dimethyl-1H-pyrrol-1-yl) benzoic acid | RmlB + RmlC | 2.67 |

### Ri03 prevents growth of other pathogenic streptococci

We extended our experiments to include other human and animal streptococcal species for which dTDP-rhamnose is an essential nucleotide sugar. GCS contains a related surface carbohydrate, the characteristic group C carbohydrate, composed of a polyrhamnose backbone decorated with a di-GalNAc side chain (Mistou *et al.*, 2016). Ri03 inhibited growth of GCS with an IC_50_ of 0.42 mM (Fig. 6B) and of *S. mutans* with an IC_50_ of 0.41 mM (Fig. 6B).

## Discussion

In this study we show that GAS RmlB and RmlC are the dTDP-glucose-4,6-dehydratase and dTDP-4-keto-6-deoxyglucose-3,5-epimerase enzymes, respectively. Both GAS RmlB and RmlC functionally replaced *S. mutans* homologs in a heterologous expression system. The rescue of growth and morphological phenotypes of the *rmlB* and *rmlC* mutants by plasmid overexpression excludes the occurrence of polar effects within the *rmlACB* operon. In addition to proving the function of both GAS RmlB and RmlC as rhamnose biosynthesis enzymes, we also identified Y159 in RmlB and H76 and K82 in RmlC as critical catalytic residues. Our complementation studies indicate that the mechanism of catalysis is conserved among streptococcal species.

Deletion of *rmlB* and *rmlC* in *S. mutans* resulted in a phenotype similar as previously observed for an *rmlD* deletion mutant (van der Beek *et al.*, 2015), underscoring that inhibition of the dTDP-rhamnose biosynthesis pathway severely impacts bacterial viability. Indeed, the *rmlABCD* genes are essential for *S. mutans* in the competitive environment of a mutant transposon library (Shields *et al.*, 2018), similar to GAS and GCS (Charbonneau *et al.*, 2017, Le Breton *et al.*, 2017, Le Breton *et al.*, 2015). We are however able to construct *rml* deletion mutants in *S. mutans* in isolation in contrast to similar attempts in GAS (van Sorge *et al.*, 2014). The reason for this discrepancy is currently unclear but may be caused by differential compositions of the cell walls or possibly these deletions are partially rescued through interacting pathways.

Screening a chemical compound library to identify inhibitors of the dTDP-L-rhamnose biosynthesis pathway is methodically challenging. Previously, indirect methods were used to determine the production of the end product dTDP-L-rhamnose by monitoring the production of cofactor NAD(P)H (Graninger *et al.*, 1999, Sivendran *et al.*, 2010, van der Beek *et al.*, 2015). A superior assay would involve a HPLC or mass-spectrometry approach where every single step of product formation and substrate consumption could be monitored. However, it is very challenging to develop this method suitable for medium or high throughput screening. We therefore investigated an approach using recombinant enzymes in their natural state, i.e. with bound co-factors in their active site. RmlB enzymes require the co-factors NAD(P), which appears to be tightly bound in the active site as it was co-purified during RmlB crystallizations (Allard *et al.*, 2002, Beis *et al.*, 2003). This is in agreement with the regeneration of NAD during dTDP-4-keto-6-deoxy-D-glucose synthesis (Allard *et al.*, 2002). GacA/RmlD enzymes also require NADPH as a co-factor, which appears to be less tightly bound as it was not observed in the GAS GacA and *S. suis* RmlD crystal structures and therefore should be regenerated by a different enzyme (Blankenfeldt *et al.*, 2002, van der Beek *et al.*, 2015). Using the binding approach, we identified seven potential inhibitors that interacted with at least one enzyme of the dTDP-L-rhamnose biosynthesis pathway with low millimolar to high micromolar affinities. One of the most potent chemical fragments Ri03, 5-(4-chlorophenyl)-2-furoic acid, is a small molecule, which binds RmlB and GacA in the high μM range, and RmlC in the low millimolar range. Despite the fact that these are only fragments, which are generally known to have low affinity and limited specificity, we demonstrated that Ri03 prevented growth of several streptococcal strains, which express rhamnose as an essential component of their cell wall. The high sequence identity of RmlB and GacA/RmlD amongst GAS, GCS and *S. mutans* (92-94% and 82-87% protein sequence identity, respectively; Fig. 2C) supports the observation that Ri03 shows similar IC_50_ values against all three strains.

Future experiments aim to investigate the binding mode of Ri03 on a structural level to the proteins RmlB, RmlC and GacA to elucidate whether Ri03 inhibits RmlB, RmlC and GacA by competing with the nucleotide-sugar and if a common binding mode is observed amongst the different enzymes. With more functional and structural knowledge, Ri03 can be further optimized to identify more potent and specific derivatives with greater potency targeting the GAS and related homologous Rml enzymes in other pathogenic species.

## Experimental procedures

### Bacterial strains and growth conditions

Bacterial strains used in this study are described in Table 2. *S. mutans* Xc (Koga *et al.*, 1989), a serotype c wild-type (WT) strain, was a kind gift of Dr. Y. Yamashita (Kyushu University, Japan) and was routinely grown in Todd-Hewitt Broth (THB, Oxoid) or on THB agar at 37°C with 5% CO_2_. When appropriate, *S. mutans* was cultured with 10 μg/ml erythromycin (ERY) or 3 μg/ml chloramphenicol (CHL). GAS 5448, a representative of the epidemic M1T1 clone, was cultured in Todd-Hewitt Broth (Becton Dickinson) supplemented with 1% yeast extract (THY; Oxoid) or on THY agar at 37°C. *E. coli* MC1061 was used for cloning purposes and was grown in lysogeny broth (LB, Oxoid) or on LB agar containing 10 μg/ml CHL at 37 °C.

**Table 2.** Bacterial strains used in this study.

| Bacterial strains | Abbreviation | Resistance |
| --- | --- | --- |
| *S. mutans* Xc wild type (serotype c) | SMU WT | - |
| *S. mutans* Xc ΔrmlB | SMU ΔrmlB | ERY |
| *S. mutans* Xc ΔrmlB + pDC123_SMU_rmlB | SMU ΔrmlB + pSB | ERY + CHL |
| *S. mutans* Xc ΔrmlB + pDC123_GAS_rmlB | SMU ΔrmlB + pGB | ERY + CHL |
| *S. mutans* Xc ΔrmlB + pDC123_GAS_rmlB_Y159F | SMU ΔrmlB + pGB_Y159F | ERY + CHL |
| *S. mutans* Xc ΔrmlC | SMU ΔrmlC | ERY |
| *S. mutans* Xc ΔrmlC + pDC123_SMU_rmlC | SMU ΔrmlC + pSC | ERY + CHL |
| *S. mutans* Xc ΔrmlC + pDC123_GAS_rmlC | SMU ΔrmlC + pGC | ERY + CHL |
| *S. mutans* Xc ΔrmlC + pDC123_GAS_rmlC_H76N | SMU ΔrmlC + pGC_H76N | ERY + CHL |
| *S. mutans* Xc ΔrmlC + pDC123_GAS_rmlC_K82A | SMU ΔrmlC + pGC_K82A | ERY + CHL |
| *S. pyogenes* 5448 wild type (serotype M1T1) | GAS | - |
| *S. equi* subsp. *zooepidemicus* MGCS10565 | GCS | - |
| *E. coli* MC1061 | - | - |
| *E. coli* BL21 (DE3) | - | - |
| *E. coli* DH5alpha | - | - |

### Cloning, expression and purification of GAS enzymes

GAS RmlB, RmlC and GacA proteins were produced and purified as described previously (van der Beek *et al.*, 2015).

### *Genetic manipulation of* S. mutans

*S. mutans* is naturally competent and was transformed as described previously. Shortly, bacteria were grown in THB containing 5% heat-inactivated horse serum (BioRad) and supplied with 500 ng knockout construct or complementation plasmid (van der Beek *et al.*, 2015). Cultures were plated on THB agar plates containing the appropriate antibiotics. Single colonies were selected and verified for the deletion of *rmlB/rmlC* and/or the presence of complementation plasmid using colony PCR and sequencing.

Deletion mutants of *rmlB* (SMU Δ*rmlB*) and *rmlC* (SMU Δ*rmlC*) were obtained via the addition of a knockout construct to competent *S. mutans* WT. This construct consisted of an ERY resistance cassette with ^~^700 bp flanking regions homologous to the up- and downstream regions of *rmlB* or *rmlC*. A detailed cloning strategy is available in the supplemental material and Table S1.

### S. mutans *complementation plasmids*

*S. mutans rmlB/rmlC* and GAS *rmlB/rmlC* were amplified from gDNA using primers containing Xbal, Xhol or BamHI restriction site (Table S1). Digested PCR products were subsequently ligated into expression plasmid pDC123 and propagated in *E. coli* MC1061 to obtain large quantities of complementation plasmid. Point mutations were introduced by mutagenesis PCR, using overlapping primers with the corresponding mutation site integrated. The constructed plasmids with the single point mutations were all confirmed by DNA sequencing.

SMU Δ*rmlB* and SMU Δ*rmlC* knockout mutants were not transformable likely resulting from the severe growth defects. Therefore, SMU Δ*rmlB* and SMU Δ*rmlC* strains complemented with *S. mutans rmlB/rmlC* or GAS *rmlB/rmlC* were constructed using a two-step approach. First, *S. mutans* WT was transformed with the complementation plasmid as described above. Next, these complemented strains were transformed with the respective *rmlB* or *rmlC* knockout constructs and selected for double antibiotic resistance.

### Scanning Electron Microscopy

Scanning electron microscopy was performed as described previously by van der Beek et al (van der Beek *et al.*, 2015). In short, bacteria were grown to mid-exponential phase, except when incubated with rhamnose inhibitor. In this case, bacteria were cultured as described below in bacterial growth inhibition assays and grown overnight. Bacteria were subsequently washed, fixed, dehydrated, mounted onto 12.5 mm specimen stubs (Agar scientific, Stansted, Essex, UK) and coated with gold to 1 nm using a Quorum Q150R S sputter coater at 20 mA. Samples were visualized with a Phenom PRO desktop scanning electron microscope (Phenom-World BV) at an acceleration voltage of 10 kV.

### S. mutans *growth curves*

Overnight cultures of *S. mutans* strains with an optical density at 600 nm (OD_600_) higher than 0.35 were diluted 10 times and grown for 1.5 h to early exponential phase. Cultures were then diluted to OD_600_ 0.03 and OD_600_ was measured manually every 30 min. For cultures with an overnight OD_600_ below 0.35, the initial 10-fold dilution and growth step was omitted. Instead, such overnight cultures were directly diluted to OD_600_ 0.03 and OD_600_ was measured every hour. All cultures were incubated at 37 °C with 5% CO_2_ in between measurements.

### Cell wall carbohydrate composition analysis

Cell wall polysaccharides were isolated from *S. mutans*, hydrolyzed and analyzed by chromatography as described previously by van der Beek *et al.* (van der Beek *et al.*, 2015). In short, bacterial cells were harvested from 2-5 L cultures and disrupted in 0.1 M citrate buffer (pH 4.6) using a bead beater (Biospec). Cell walls were collected by centrifugation and boiled for 1 h at 100 °C in 0.1 M sodium acetate (pH 4.6) containing 4% sodium dodecyl sulfate. Samples were subsequently treated with RNase, DNase, pronase E and trypsin. Cell walls containing peptidoglycan and the serotype C carbohydrate were lyophilized before hydrolysis with TFA. Finally, carbohydrate analysis of monosaccharides was performed on a Dionex ICS-3000 Ion Chromatography System (Dionex / Thermo Scientific, 1228 Titan Way, P.O. Box 3603, Sunnyvale, CA, 94088-3603, United States) using a CarboPac PA20 (Dionex / Thermo Scientific, 1228 Titan Way, P.O. Box 3603, Sunnyvale, CA, 94088-3603, United States) 3 × 150 mm column, equipped with a ICS-3000 Electrochemical Detector (Dionex / Thermo Scientific, 1228 Titan Way, P.O. Box 3603, Sunnyvale, CA, 94088-3603, United States). Monosaccharides from all samples were eluted from the column with 12% 0.2 M NaOH, except for monosaccharides from SMU Δ*rmlB* and Δ*rmlC*, which were eluted with 8% 0.2 M NaOH. The ‘Carbohydrates (Standard Quad)’ waveform was used for detection.

### BLI screen using recombinant RmlB, RmlC and GacA

The Maybridge library of fragment-like molecules (Ro3) was purchased from Maybridge (USA). All three GAS proteins, RmlB, RmlC and GacA, were screened against the library using identical protocols. Proteins were biotinylated on primary amine amino acid side chains using the Thermo Scientific™ EZ-Link™ NHS-PEG4-Biotin reagent. The enzymes were incubated with NHS-PEG_4_-Biotin in a 1:1 molar ratio at 150 μM for 45 min at room temperature. The reaction buffer contained 25 mM HEPES, pH 7.5, 150 mM NaCl and 0.02 mM TCEP. Excess NHS-PEG_4_-biotin reagent was removed using 2 mL Zeba desalting spin columns (Thermo Scientific). Successful biotinylation on primary amines was investigated via Western blotting and probing with ExtrAvidin^®^–Peroxidase antibody, 1:10,000 (Sigma E2886, Fig. S1A). Before compound screening, the proteins were loaded onto a parallel set of superstreptavidin biosensors (SSA) by incubation in buffer for 900 sec. Protein concentrations suitable for the experiments were determined in a series of dilutions. Final screening was conducted at concentrations of: 50 μg/mL RmlB, 25 μg/mL RmlC and 12.5 μg/mL GacA. All compounds were at 2% DMSO concentration. The sensor was blocked by immersion in biocytin (10 μg/mL) for 30 sec.

Initial library screening was performed at a compound concentration of 200 μM. Hits were identified by plotting the response rate of every single compound after background subtraction. In total, 12 compounds were validated in 6-point concentration series using 3-fold dilutions, ranging from 670 μM to 3 μM. Data were processed and kinetic parameters were calculate using the ForteBio software. Binding curves were manually inspected and approximate binding constants K_D_ are listed in Table S3.

### Bactericidal activity

GAS 5448, GCS MGCS10565 and *S. mutans* Xc strains (Table 2) were grown in appropriate bacterial culture broth overnight at 37 °C. Bacterial cultures were diluted 10 times and grown to mid-log phase (OD_600_ 0.4). Cultures were diluted 200 times in culture medium followed by a 2-fold dilution with various concentrations of rhamnose inhibitor or DMSO. OD_600_ was recorded every 15 min at 37°C with 5 sec medium shaking before measurement for GAS, GCS and *S. mutans* using either a Bioscreen C MBR machine (Growth Curves AB Ltd, Oy, Finland) or a Synergy 2 plate reader (Biotek). OD_600_ values were plotted against the concentration of compound after 9 hours of growth using GraphPad Prism 7. IC_50_ values were calculated using a non-linear four-parameter dose-response curve with variable slope. Because high concentrations were not always feasible due to solubility issues, a high concentration of compound (50 mM) was artificially introduced where appropriate and set to zero to obtain more reliable dose-response curves.

## Acknowledgements

The authors would like to acknowledge Dr Olawale Raimi (University of Dundee) for assistance with BLI. We are grateful for funding to HCD and his laboratory from Tenovus Scotland, Wellcome Trust and Royal Society (grant number 109357/Z/15/Z.), FEBS and the University of Dundee. This work was supported by a VIDI grant (91713303) from the Dutch Scientific Organization (NWO) to N.M.v.S and S.v.d.B. The authors declare no conflict of interest.

## Author contributions

Conception and design of the study: SvdB, AZ, HCD, NvS; Acquisition, analysis and interpretation of the data: SvdB, AZ, EC, RNC, BHM, GJB, HCD, NvS; Writing of the manuscript: SvdB, AZ, HCD, NvS

## Abbreviated summary

Biosynthesis of the nucleotide sugar precursor dTDP-L-rhamnose through the enzymes RmlABCD is critical for the viability and virulence of many human pathogenic bacteria. We confirm the function and the catalytic residues of *Streptococcus pyogenes* RmlB and RmlC through bacterial genetics and carbohydrate analysis. Screening 1,000 compounds against recombinant enzymes, we identified a compound Ri03 that affect streptococcal viability and can serve as a starting point for development of new inhibitors targeting the RmlABCD pathway.

---

**Fig. S1.**
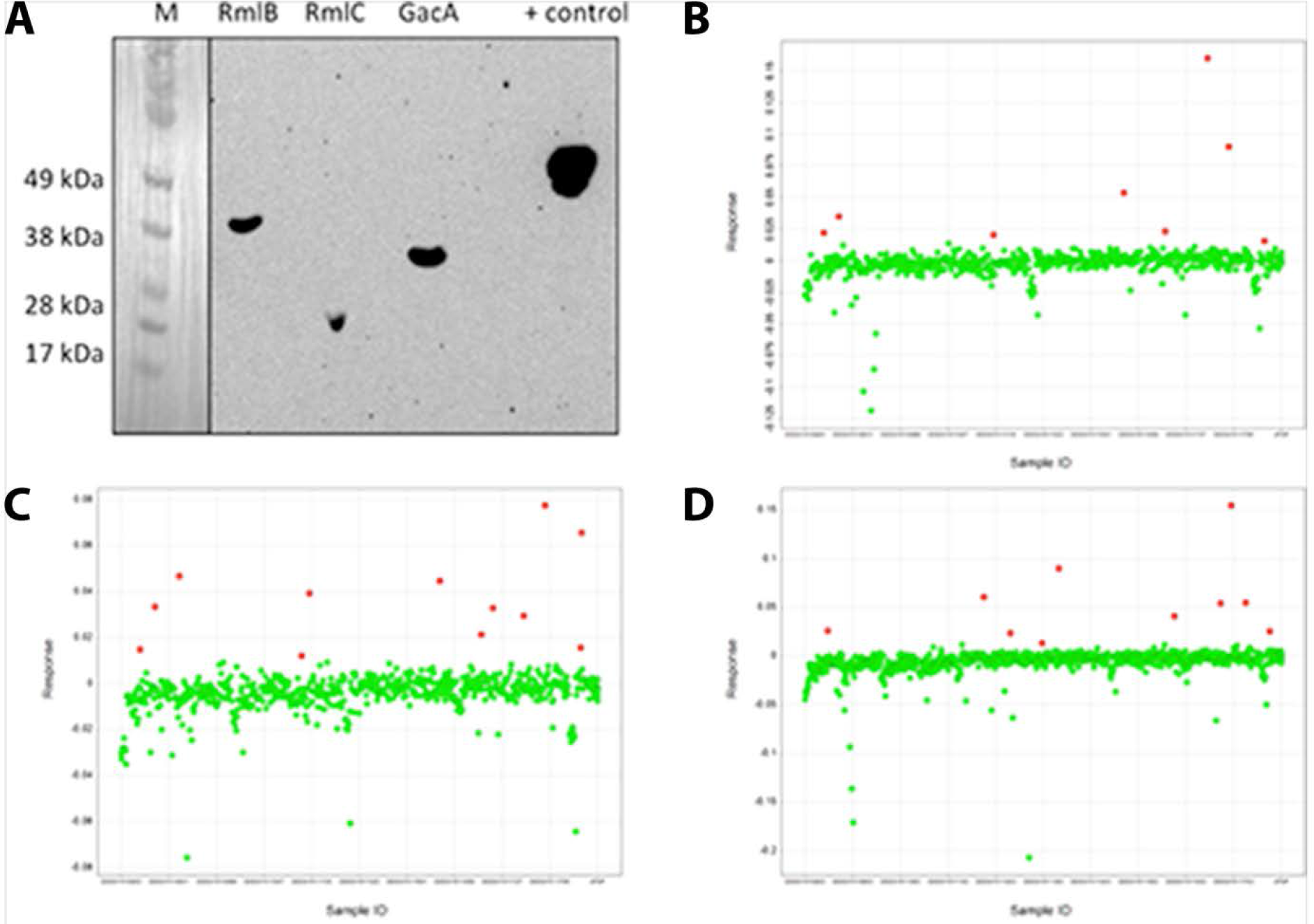
Mayflower library screening data. (A) Anti-biotin Western blot of purified RmlB, RmlC and GacA proteins (RmlB = 39 kDa, RmlC = 23 kDa, GaCA = 32 kDa), with positive control (biotinylated Protein X; 55 kDa). (B) RmlB, (C) RmlC and (D) GacA hit cut-off value figures from BLI screen. The binding curve of all compounds were analyzed with robust calculations and plotted as compound (X-axis) vs. response. Potential hits above the hit cut-off are depicted in red. In total 17 compounds were selected and tested in dose-dependent BLI studies against all three target enzymes.

**Fig. S2.**
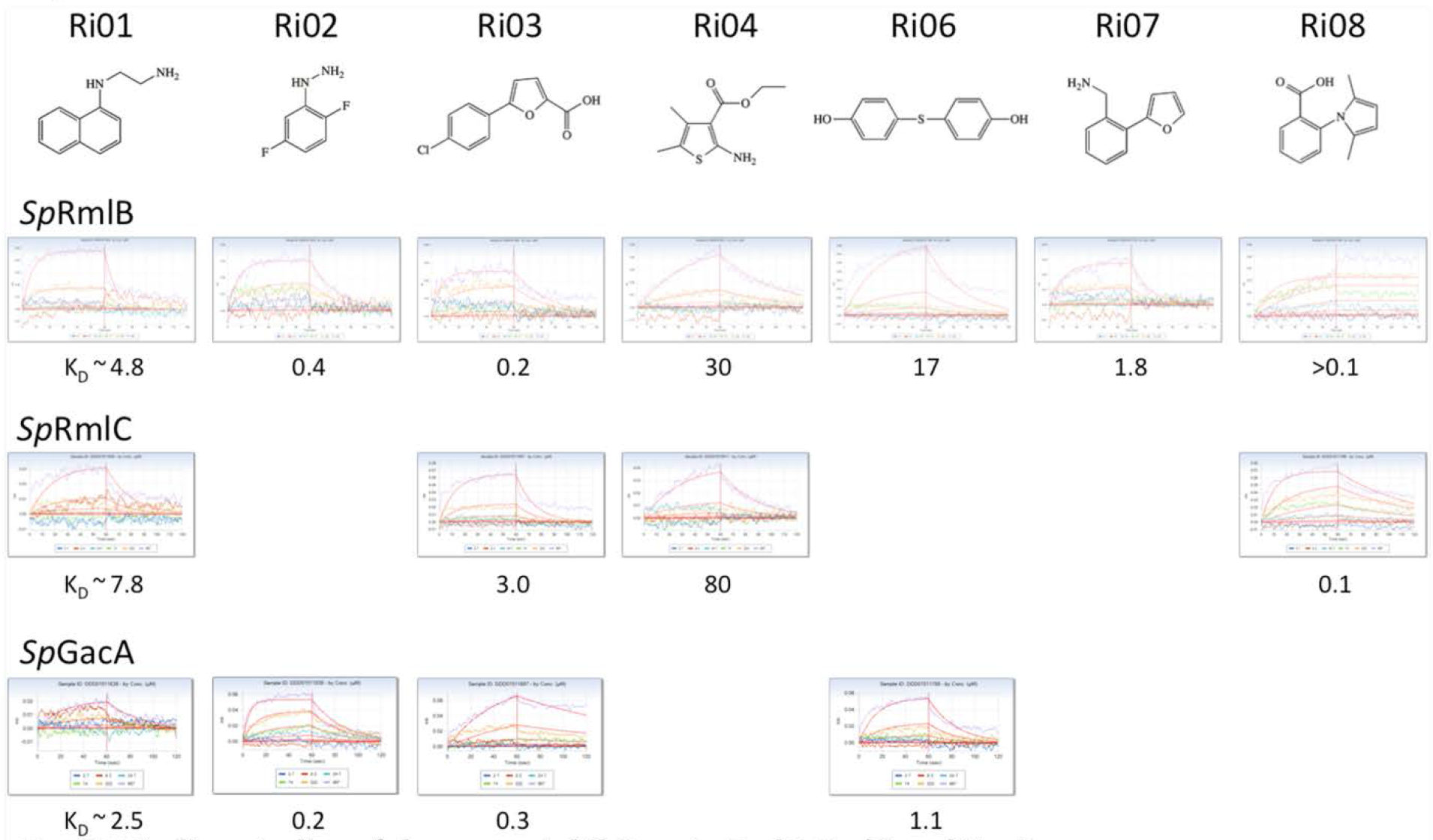
Binding studies of rhamnose inhibitors to RmlB, RmlC and GacA. Chemical structures of identified inhibitors, binding curves vs. purified recombinant proteins and calculated binding affinities (K_D_) in mM.

